# X-ray crystallographic analyses of 14 IPMK inhibitor complexes

**DOI:** 10.1101/2024.05.09.593385

**Authors:** Huanchen Wang, Raymond D. Blind, Stephen B. Shears

## Abstract

Inositol polyphosphate multikinase (IPMK) is a ubiquitously expressed kinase that has been linked to several cancers. Here, we report 14 new co-crystal structures (1.7Å - 2.0Å resolution) of human IPMK complexed with various IPMK inhibitors developed by another group. The new structures reveal two ordered water molecules that participate in hydrogen-bonding networks, and an unoccupied pocket in the ATP-binding site of human IPMK. New Protein Data Bank (PDB) codes of these IPMK crystal structures are: **8V6W**(1.95Å), **8V6X**(1.75Å), **8V6Y**(1.70Å), **8V6Z**(1.85Å), **8V70**(1.85Å), **8V71**(1.70Å), **8V72**(2.0Å), **8V73**(1.90Å), **8V74**(1.85Å), **8V75**(1.85Å), **8V76**(1.95Å),**8V77**(1.95Å), **8V78**(1.95Å), **8V79**(1.95Å).

## Introduction

Inositol polyphosphate multikinase (IPMK) is a ubiquitously expressed kinase, the first crystal structure of any IPMK orthologue published was that of the yeast ipk2(11), followed by the crystal structure of the plant *Arabidopsis thaliana* orthologue(12). These crystal structures established IPMK has the typical kinase fold, containing an N-terminal lobe and C-terminal lobe with the ATP binding site at the interface between these two lobes. The structure of the plant IPMK showed a larger IP-binding loop, suggesting why the plant orthologue has no activity on the phospholipid PI(4,5)P2(7), yet retains inositol phosphate kinase activity(12). X-ray crystal structures of the human IPMK kinase domain were solved in close succession by two independent labs, in the apo form(13) and in complex with nucleotide(14), and with the kinase substrates inositol phosphate and the phosphoinositide lipid PI(4,5)P2(14). These structures and kinetic analyses revealed several interesting aspects of IPMK catalysis and regulation. However, no inhibitor-bound structures of IPMK had been solved to that point.

X-ray structural analyses of flavonoids bound to human IPMK revealed hydrophobic and polar interactions between the flavonoids and particular amino acid side chains in IPMK(15). However these ATP-competitive flavonoids are inhibit many diverse kinases. These studies also suggested that ordered water molecules in the ATP-binding site might be important for flavonoid interaction with IPMK, and informed potential inhibiting properties in the development of inositol-phosphate kinase inhibitors. However, flavonoids can more generally inhibit protein and phospholipid kinase activity, so structure-based improvements would likely require improvements to flavonoid interactions with human IPMK. Thus it would be useful if new chemicals could be developed that were specific to IPMK. In the process of identifying IP6K1 inhibitory molecules (16, 17) some molecules were found to inhibit IPMK, which were developed by another group and will be reported elsewhere. Here, we report 14 new X-ray crystal structures of the IPMK kinase domain complexed with IPMK inhibitors.

## Results

### PDB coordinates of 14-inhibitor bound IPMK crystal structures

We solved 14 crystal structures of inhibitors complexed with the human IPMK kinase domain **8V6W**(1.95Å), **8V6X**(1.75Å), **8V6Y**(1.70Å), **8V6Z**(1.85Å), **8V70**(1.85Å), **8V71**(1.70Å), **8V72**(2.0Å), **8V73**(1.90Å), **8V74**(1.85Å), **8V75**(1.85Å), **8V76**(1.95Å), **8V77**(1.95Å), **8V78**(1.95Å), **8V79**(1.95Å). The 1.95Å X-ray crystal structure complex of **PDB:8V6W** confirms what has been established in other crystallographic studies of IPMK(15), the nucleotide-binding site of human IPMK adopts the typical kinase N-lobe and C-lobe fold connected by a hinge loop and the nucleotide binding site occupied by an ambiguous electron density interpreted as inhibitor. Four hydrogen bonds were formed with the IPMK hinge region, including three hydrogen bonds with IPMK backbone residues and one hydrogen bond with Asp132. For comparison, the adenine group of ATP in the co-crystal structure with IPMK only makes 2 hydrogen bonds with the polypeptide backbone(14).

The overall fold of the IPMK kinase domain is similar to those structures already published in complex with various other small molecules and substrates. **PDB:8V6W** also shows four polar amino acids in the IPMK back-pocket (Lys75, Glu86, Tyr90, and Asp385) formed a hydrogen bond network with the small molecule. The residues Lys75, Glu 86, and Asp385 comprise the catalytic triad in IPMK, these residues are conserved in inositol phosphate kinases (13, 19). However, Tyr90 is unique to IPMK and IP6K, and is not present in IP3K or the related PKA protein kinase. Thus, we named residues Lys75, Glu86 and Tyr90 the “KEY motif”, with this motif specific to inositol phosphate kinases. **PDB:8V6W** shows IPMK anchored the small molecules with two hydrogen bond clusters, suggesting a two-point anchoring recognition mechanism, forming hydrophobic interactions with residues Ile65, Val73, P111 and Leu254 of IPMK, also observed in the complex of ATP with IPMK. Thus, the new 1.95Å X-ray crystal structure complex of **PDB:8V6W** reveals multiple structure-based elements that can be used in future analyses of IPMK.

The 1.75Å X-ray crystal structure of **PDB:8V6X**, the 1.70Å structure of **PDB:8V6Y** and the 1.85Å crystal structure of **PDB:8V6Z** all show a well-coordinated water we call “water1”, which can have a variable number of hydrogen bonding interactions with small molecules. The 1.85Å structure of **PDB:8V6Z** suggested another water2 molecule with less well-defined density. Further, the 1.85Å structure of **PDB: 8V70**, the 1.70Å structure of **PDB: 8V71**, and the 2.0Å structure of **PDB: 8V72** all show a coordinated water2 molecule in the active site, similar to structures of the 1.90Å **PDB: 8V73** and 1.90Å, **PDB: 8V74**. However, the 1.85Å structure of **PDB:8V75** showed water2 and a new water3 molecule in the active site, forming a hydrogen bond network with Arg182, while the 1.9Å structure of **PDB:8V76** still had water2 and water 3 ordered but did not show interaction with the same Arg182. The additional water3 molecule, together with water2, forms an internal hydrogen bond network. The 1.95Å crystal structure of **PDB: 8V77** was less revealing that the 1.95Å structure of **PDB: 8V78**, showing two polar contacts with the backbone of Gly180 and Met181. The 1.95Å crystal structure of **PDB: 8V79** shows water2 forming an internal hydrogen bond network. Together, these structures of IPMK have highlighted the importance of ordered waters in the ATP-binding active site of IPMK.

## Discussion

The new X-ray crystal structures reported here have led to some observations that can be applicable more broadly to the field of kinase structure. The hinge region, along with the C-spine, plays a significant role in inhibitor binding affinity in most protein kinases, as well as in inositol phosphate kinases(15). Generally, kinase inhibitors solely targeting the hinge region may lack selectivity, which can be achieved by targeting the back pocket. However, designing an inhibitor with back pocket binding presents additional challenges as has been previous noted(20, 21). The back pocket is situated directly behind the adenine of ATP, which is also known as the hydrophobic pocket in protein kinases. In IPMK, the side chain of the gatekeeper Leu130 is relatively shorter than a methionine residue, allowing for more variation of moiety A in the back pocket(14). Water-mediated intramolecular interactions between moieties A and B may also affect IPMK binding properties. The previously published crystal structure of quercetin bound to IPMK showed a similar ordered water in the crystal structure(15), indicating the potential importance of ordered water to IPMK ligand binding and selectivity. Importantly, in that study several other structures solved complexed with natural product flavonoids did not have a water molecule ordered in those structures(15). In the crystal structures here, ordered water molecules were observed in the active site, but some structures showed hydrogen bonding networks suggesting these networks may be important. The role of ordered waters highlights the power of X-ray crystallography in revealing binding mechanism, highlight the value of high-resolution crystallography in studying small molecule binding to enzymes, such as IPMK.

## Acknowledgements

This work was supported by Intramural Research Program grant of the National Institute of Environmental Health Sciences to S.B.S., R21 AG07197 and National Institutes of General Medical Sciences (NIGMS) R01 GM132592 to R.D.B.

## Materials and Methods

### Xray crystallography structural studies

Recombinant human IPMK kinase domain was prepared as previously described (18). The purity was estimated to be >80% as judged by SDS-PAGE. The purified protein was stored in aliquots at −80°C. Crystals of human apo-IPMK were prepared as described previously. Complex crystals were produced by soaking apo-crystals into a mixture of 2– 10 mM compounds with 35% (w/v) PEG 400, 0.1 M Li2SO4, 100 mM HEPES (pH 7.5) at 25°C for 3 days. Diffraction data were collected using APS beamlines 22-ID and 22-BM. All data were processed with the program HKL2000. The crystal structures were determined by using rigid body and direct Fourier synthesis, and refined with the equivalent and expanded test sets by using programs in the CCP4 package. The molecular graphics representations were prepared with the program PyMol (Schrödinger, LLC). Atomic coordinates and structure factors for human IPMK/compound co-complexes have been deposited with the Protein Data Bank with the following accession codes 8V6W, 8V6X, 8V6Y, 8V6Z, 8V70, 8V71, 8V72, 8V73, 8V74, 8V75, 8V76, 8V77, 8V78, 8V79, structural refinement statistics are shown in Table 1.

**Table 1.**
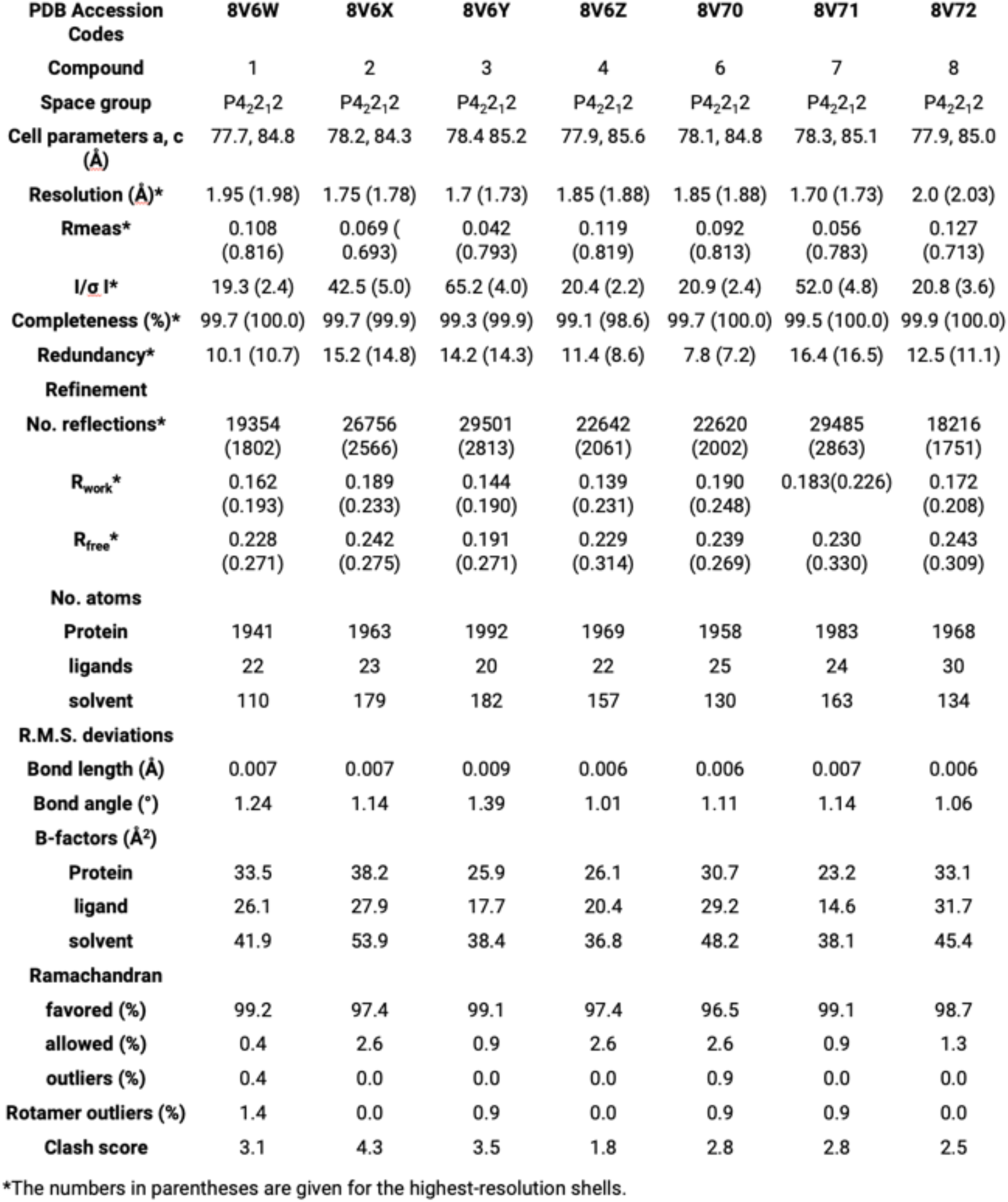
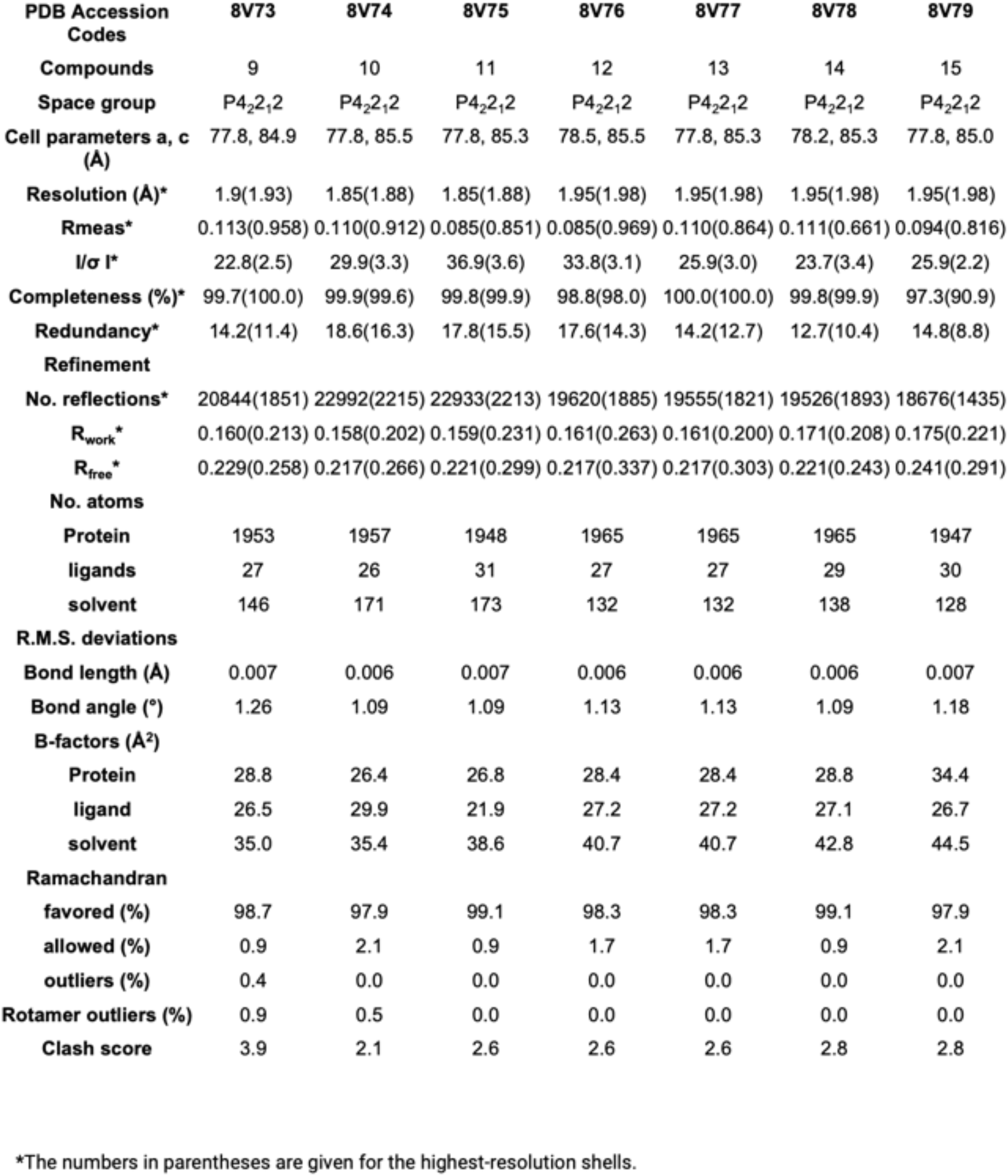
Data collection and refinement statistics.

## Notes

### Competing Interest Statement

The authors have declared no competing interest.

